# Deep transfer learning provides a *Pareto* improvement for multi-ancestral clinico-genomic prediction of diseases

**DOI:** 10.1101/2022.09.22.509055

**Authors:** Yan Gao, Yan Cui

## Abstract

Accurate genomic predisposition assessment is essential for the prevention and early detection of diseases. Polygenic scores and machine learning models have been developed for disease prediction based on genetic variants and other risk factors. However, over 80% of existing genomic data were acquired from individuals of European descent. As a result, clinico-genomic risk prediction is less accurate for non-European populations. Here we employ a transfer learning strategy to improve the clinico-genomic prediction of disease occurrence for the data-disadvantaged populations. Our multi-ancestral machine learning experiments on clinico-genomic datasets of cancers and Alzheimer’s disease and synthetic datasets with built-in data inequality and subpopulation shift show that transfer learning can significantly improve disease prediction accuracy for data-disadvantaged populations. Under the transfer learning scheme, the prediction accuracy for the data-disadvantaged populations can be improved without compromising the prediction accuracy for other populations. Therefore, transfer learning provides a *Pareto* improvement toward equitable machine learning for genomic medicine.

## Introduction

Predicting disease occurrence based on clinico-genomic data is important to precision medicine. Traditionally, disease prediction was primarily based on epidemiological risk factors such as lifestyle variables and family history. The recent advances in high-throughput genotyping and genome sequencing technologies have enabled genome-wide association studies (GWAS) of large cohorts. The data and discoveries from GWAS provide a foundation for polygenic risk prediction for complex diseases. However, over 80% of the existing GWAS data were acquired from people of European descent^1-7^, and the ancestral (or ethnic) diversity in GWAS has not improved in recent years^1,6^. The genomic data paucity for the non-European populations, which constitute about 84% of the world’s population, leads to low-quality artificial intelligence (AI) models for these data-disadvantaged populations. Genomic data inequality has become a health hazard for the vast majority of the world’s population and a new source of health disparities^8^.

The polygenic scores used for disease risk assessment^9-16^ are often inadequate for accurate individualized disease prediction. This is partially because 1) complex diseases are caused not only by genetic factors but by a combination of genetic, environmental, and lifestyle factors; and 2) linear polygenic scores lack the expressive power and model capacity to capture the non-linear, non-additive interactions inherent in the complex genotype-phenotype relationship. Non-additive genetic interactions can significantly reduce the accuracy of polygenic prediction^17^. Recently, deep learning and other machine learning models were developed to integrate genomic and epidemiological factors for disease prediction and prognosis^18-20^. The machine learning models that excel at capturing complex non-linear interactions generally outperform conventional disease prediction models^20-23^.

Polygenic scores derived from GWAS of predominantly European ancestry generalize poorly to other ancestry groups^6,24-29^. Recent studies show that the polygenic prediction for non-European populations can be improved by calibrating parameters for genetic effect sizes or model sparsity (or shrinkage) patterns across ancestry groups in various regression models^30-35^. However, the linearity, additivity, distribution normality, and other assumptions made by these polygenic models limit their capacity to learn and transfer complex representations across ancestry groups. Recently, we developed deep neural network (DNN) based transfer learning methods to reduce the disparity in multi-ancestral (or multi-ethnic) machine learning for cancer progression and survival prognosis from mRNA and protein expression data^8,36^. In this work, we employ a deep transfer learning strategy for disease prediction from multi-ancestral GWAS data. We compare the performance of the transfer learning scheme to that of the current prevalent multi-ancestral machine learning schemes and find that transfer learning can significantly improve disease prediction for the data-disadvantaged ancestral groups.

Fairness-aware machine learning methods^37^ generally rely on various *ad hoc* constraints or penalty terms in the loss function and training process to enforce performance parity across different subpopulations, resulting in a dilemmatic fairness-accuracy tradeoff^38,39^. The transfer learning scheme is not subject to such fairness-accuracy tradeoff. The machine learning model trained for a data-abundant population can be utilized to aid in training the data-disadvantaged populations without reducing its prediction accuracy for the data-abundant population. Thus, transfer learning can provide a *Pareto* improvement^40^, in which the prediction accuracy can be improved for the data-disadvantage populations while remaining unchanged for the data-abundant populations (**Fig.1**).

**Fig. 1.**
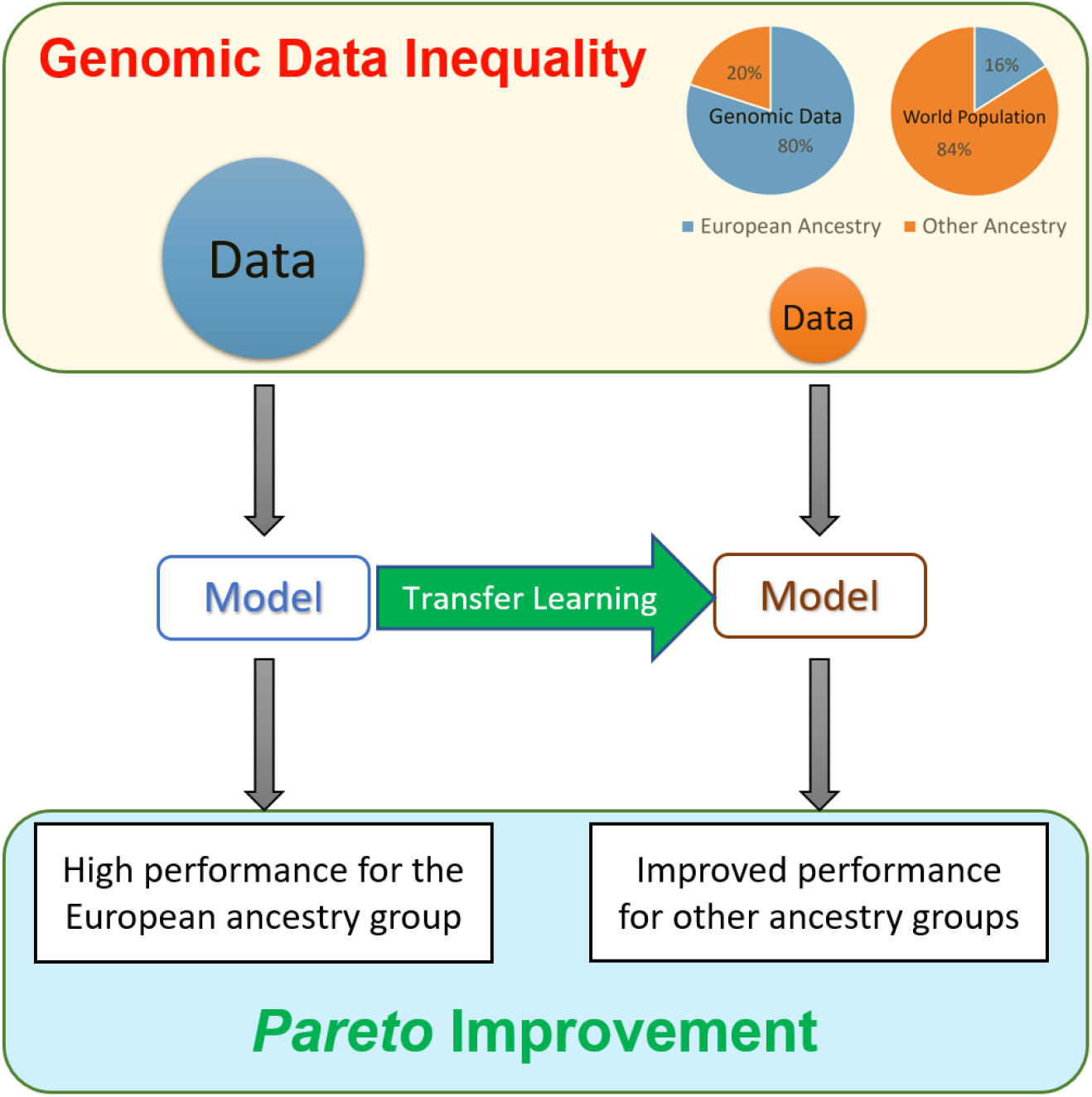
Transfer learning provides a *Pareto* improvement toward equitable machine learning for genomic medicine. The conspicuous genomic data inequality confers a significant health risk to the data-disadvantaged populations. The transfer learning scheme can improve the performance of AI models for the data-disadvantaged populations by consulting but not affecting the model for the data-rich population, thereby leading to a *Pareto* improvement.

## Results

We defined three categories of multi-ancestral (or multi-ethnic) machine learning schemes based on how the data from different subpopulations are used: mixture learning, independent learning, and transfer learning^8^ (**Table 1**). Mixture learning uses data from all ancestral groups indistinctly for model training and testing. Independent learning trains and tests a model for each ancestral group separately. In transfer learning, a model is first trained on the data of the European population (source domain), then the knowledge learned during the training is transferred to assist the model development for a DDP (target domain). Currently, the mixture and independent learning schemes are generally used for machine learning with multi-ancestral GWAS data. In our machine learning experiments, we find that both mixture and independent learning schemes generate models with low performance for the data-disadvantaged populations (DDPs). We also find that transfer learning can significantly improve machine learning model performance on the DDPs (**Table 1, Fig. 2**).

**Table 1.**
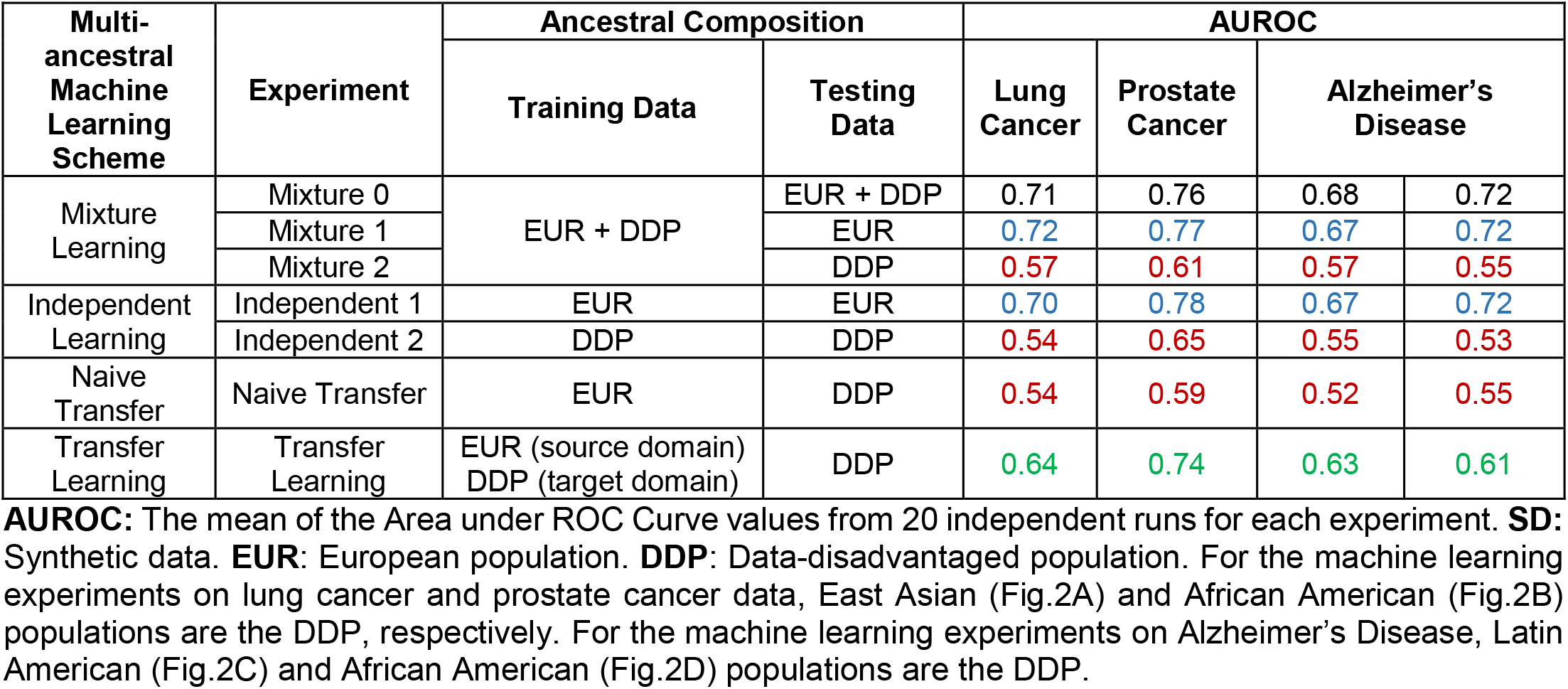
Multi-ancestral machine learning schemes and experiments.

| Multi-ancestral Machine Learning Scheme | Experiment | Ancestral Composition |  | AUROC |  |  |  |
| --- | --- | --- | --- | --- | --- | --- | --- |
|  |  | Training Data | Testing Data | Lung Cancer | Prostate Cancer | Alzheimer’s Disease |  |
| Mixture Learning | Mixture 0 | EUR + DDP | EUR + DDP | 0.71 | 0.76 | 0.68 | 0.72 |
|  | Mixture 1 |  | EUR | 0.72 | 0.77 | 0.67 | 0.72 |
|  | Mixture 2 |  | DDP | 0.57 | 0.61 | 0.57 | 0.55 |
| Independent Learning | Independent 1 | EUR | EUR | 0.70 | 0.78 | 0.67 | 0.72 |
|  | Independent 2 | DDP | DDP | 0.54 | 0.65 | 0.55 | 0.53 |
| Naive Transfer | Naive Transfer | EUR | DDP | 0.54 | 0.59 | 0.52 | 0.55 |
| Transfer Learning | Transfer Learning | EUR (source domain)<br>DDP (target domain) | DDP | 0.64 | 0.74 | 0.63 | 0.61 |

**Fig. 2.**
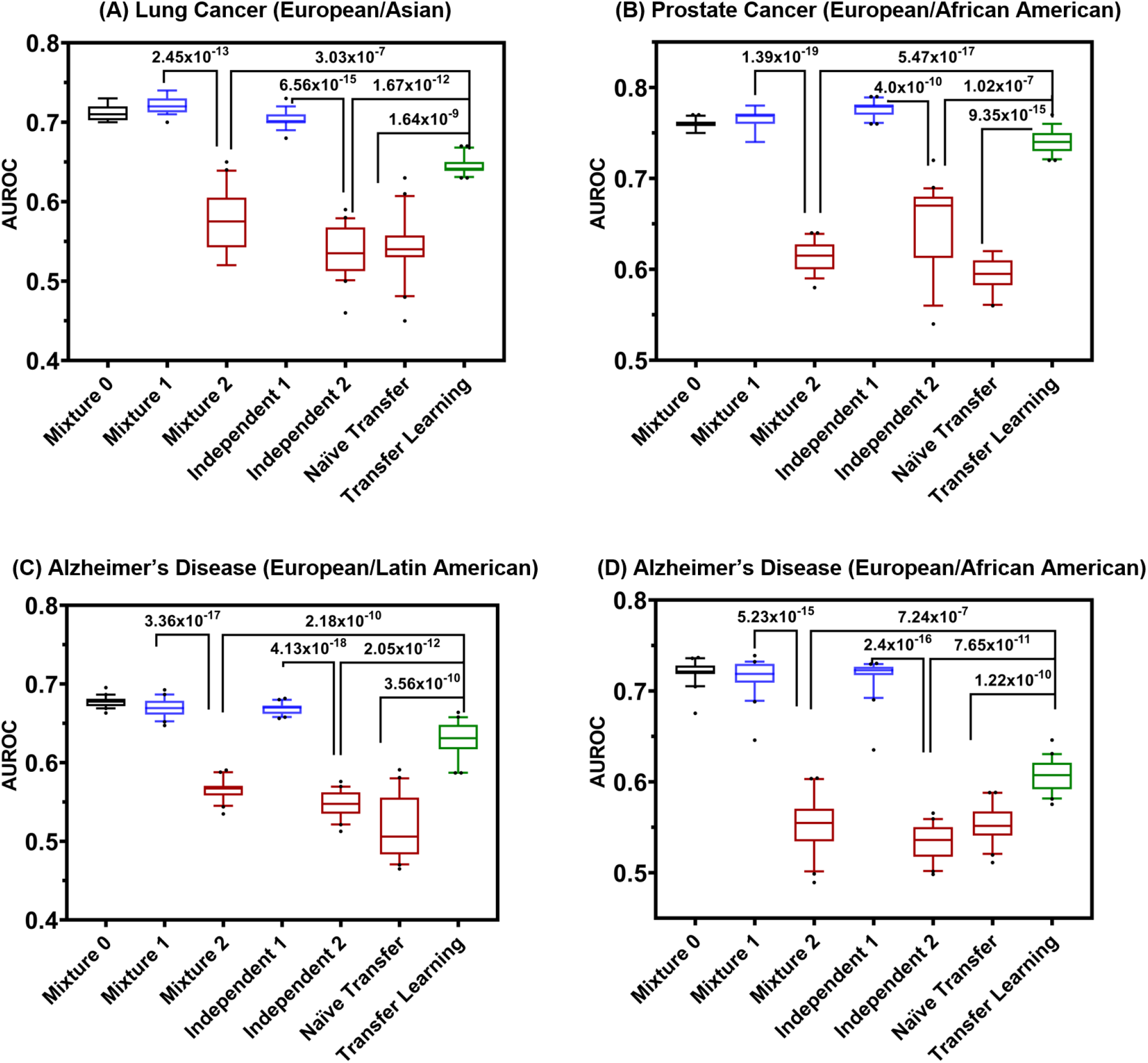
Multi-ancestral machine learning scheme comparison. Machine learning experiments for predicting case/control status for (**A**) Lung Cancer (European and East Asian), (**B**) Prostate Cancer (European and African American), (**C**) Alzheimer’s Disease (European and Latin American), and (**D**) Alzheimer’s Disease (European and African American) GWAS participants using clinico-genomic data. In each panel, the box plots show the AUROC values for the seven experiments (20 independent runs for each experiment). Box-plot elements are: center line, median; box limits, 25 and 75 percentiles; whiskers, 10–90 percentiles; points, outliers.

### Multi-ancestral machine learning experiments on GWAS data

We compared the performance of the three multi-ancestral machine learning schemes using GWAS data. We assembled the datasets for our machine learning experiments using data from multi-ancestral GWAS of lung cancer, prostate cancer, and Alzheimer’s Disease from dbGaP^41-43^, the database of Genotypes and Phenotypes. The assembled datasets for our machine learning experiments are summarized in **Table S1**.

Our machine learning tasks are to predict the disease status (case/control) from genotype and clinical data. We used the mean AUROC (Area under ROC Curve)^44^ from 20 independent runs of each machine learning experiment as the index to compare the machine learning model performance. The mixture learning scheme generated significant model performance gaps between the European and data-disadvantaged populations (**Table 1, Fig. 2**): 0.15 (*p* = 2.45 × 10^−13^) for lung cancer (European and East Asian), 0.16 (*p* = 1.39 × 10^−19^) for prostate cancer (European and African American), 0.1 (*p* = 3.36 × 10^−17^) for Alzheimer’s disease (European and Latin American), and 0.17 (*p* = 5.23 × 10^−15^) for Alzheimer’s disease (European and African American). The independent learning scheme also generated significant model performance gaps: 0.16 (*p* = 6.56 × 10^−15^) for lung cancer (European and East Asian), 0.13 (*p* = 4.0 × 10^−10^) for prostate cancer (European and African American), 0.12 (*p* = 4.13 × 10^−18^) for Alzheimer’s disease (European and Latin American), and 0.19 (*p* = 2.4 × 10^−16^) for Alzheimer’s disease (European and African American). These two schemes are widely used in multi-ancestral machine learning but are prone to generating machine learning models with low performance for data-disadvantaged populations (DDPs). Naïve transfer, in which the model trained using data from the source domain was applied directly to the target domain without adaptation, also generated models with low performance on the DDPs. The low performance of mixture learning for the DDPs (Mixture 2) and naïve transfer is consistent with previous observations of the low generalizability of polygenic scores developed using data from populations of predominantly or exclusively European ancestry. In the transfer learning scheme, a model learned from the source domain with abundant training data (the European population) was used to improve machine learning in the target domain with insufficient training data (the DDPs). In our experiments, transfer learning significantly improved the machine learning model performance (**Table 1, Fig. 2**). We define the performance disparity gap as 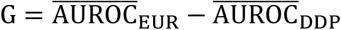, where 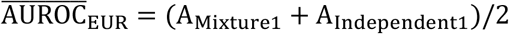, representing the average performance for the European population, and 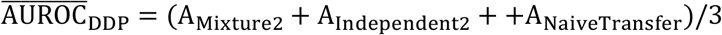, representing the average performance for the data-disadvantaged population, A_Mixture0_, A_Mixture1_, A_Mixture2_, A_Independent1_, A_Independent2_, A_NaiveTransfer_ and A_TransferLearning_ represent the mean AUROC from 20 independent runs for each of the seven experiments (**Table 1**), respectively. We observed significant model performance disparity gaps: G = 0.16 between the European and the East Asian groups in lung cancer; G = 0.158 between the European and African American groups in prostate cancer; G = 0.123 between the European and Latin American groups in Alzheimer’s disease; and G = 0.177 between the European and African American groups in Alzheimer’s disease. Transfer learning reduced the performance disparity gap by 56% for lung cancer (European and East Asian), 78% for prostate cancer (European and African American), 68% for Alzheimer’s disease (European and Latin American), and 38% for Alzheimer’s disease (European and African American).

### Multi-ancestral machine learning experiments on synthetic data

To compare the multi-ancestral machine learning schemes, we conducted experiments (as defined in **Table 1**) on the synthetic data for five ancestry groups: African (AFR), Admixed American (AMR), East Asian (EAS), European (EUR), and South Asian (SAS), representing the global genetic diversity of human populations. We downloaded the simulated genotype data of the five ancestry groups^32^ from the Harvard Dataverse^45^. We assembled four datasets to simulate the data inequality among ancestry groups. Each dataset contains genotype data of 10,000 individuals from the EUR ancestry group and 2,000 individuals from one of the four non-European ancestry groups. We randomly selected 500 SNPs as the causal SNPs for the disease. The absolute allele frequency difference for 500 causal SNPs between EUR and the other four populations is visualized using a heatmap (**Figure 3**). The heatmap shows the absolute minor allele frequency difference between the EUR population and the data-disadvantaged population populations AMR, SAS, EAS, and AFR. We constructed a statistical model to simulate the genotype-phenotype relationship (see the Methods section) and to generate the disease status (case/control) for all the individuals in the synthetic datasets. The synthetic data capture two typical characteristics of multi-ancestral datasets: the data inequality and the subpopulation shift (i.e., data distribution discrepancy between ancestry groups), which are the key factors influencing the performance of different multi-ancestral machine learning schemes^8,36^. We generated synthetic datasets using various combinations for the values of two parameters of the simulation model, *h*^2^ and *ρ*, the heritability of the disease and the correlation of the genetic effect sizes between EUR and other populations, respectively (**Table 2** and **Table S1**). The case/control ratio for all the synthetic datasets was 1:1.

**Table 2.** Multi-ancestral machine learning experiments on synthetic data.

| Synthetic Datasets |  |  |  | AUROC |  |  |  |  |  |  |
| --- | --- | --- | --- | --- | --- | --- | --- | --- | --- | --- |
| ID | DDP | $h^2$ | $\rho$ | Mixture 0 | Mixture 1 | Mixture 2 | Independent 1 | Independent 2 | Naive Transfer | Transfer Learning |
| SD1 | AMR | 0.50 | 0.80 | 0.74 | 0.75 | 0.70 | 0.76 | 0.62 | 0.62 | 0.72 |
| SD2 | SAS | 0.50 | 0.77 | 0.72 | 0.73 | 0.69 | 0.74 | 0.62 | 0.61 | 0.71 |
| SD3 | EAS | 0.50 | 0.58 | 0.73 | 0.74 | 0.67 | 0.72 | 0.64 | 0.62 | 0.69 |
| SD4 | AFR | 0.50 | 0.54 | 0.72 | 0.73 | 0.65 | 0.73 | 0.66 | 0.57 | 0.68 |
| SD5 | AMR | 0.25 | 0.80 | 0.61 | 0.62 | 0.59 | 0.62 | 0.55 | 0.55 | 0.60 |
| SD6 | SAS | 0.25 | 0.77 | 0.58 | 0.59 | 0.57 | 0.61 | 0.56 | 0.56 | 0.60 |
| SD7 | EAS | 0.25 | 0.58 | 0.57 | 0.58 | 0.55 | 0.60 | 0.56 | 0.57 | 0.59 |
| SD8 | AFR | 0.25 | 0.54 | 0.60 | 0.60 | 0.57 | 0.60 | 0.56 | 0.54 | 0.58 |

**AUROC:** The mean Area under ROC Curve values from 20 independent runs for each experiment; **SD:** Synthetic data; **EUR:** European population; **DDP:** Data-disadvantaged population; **AFR:** African; **AMR:** Admixed American; **EAS:** East Asian; **SAS:** South Asian.

**Fig. 3.**
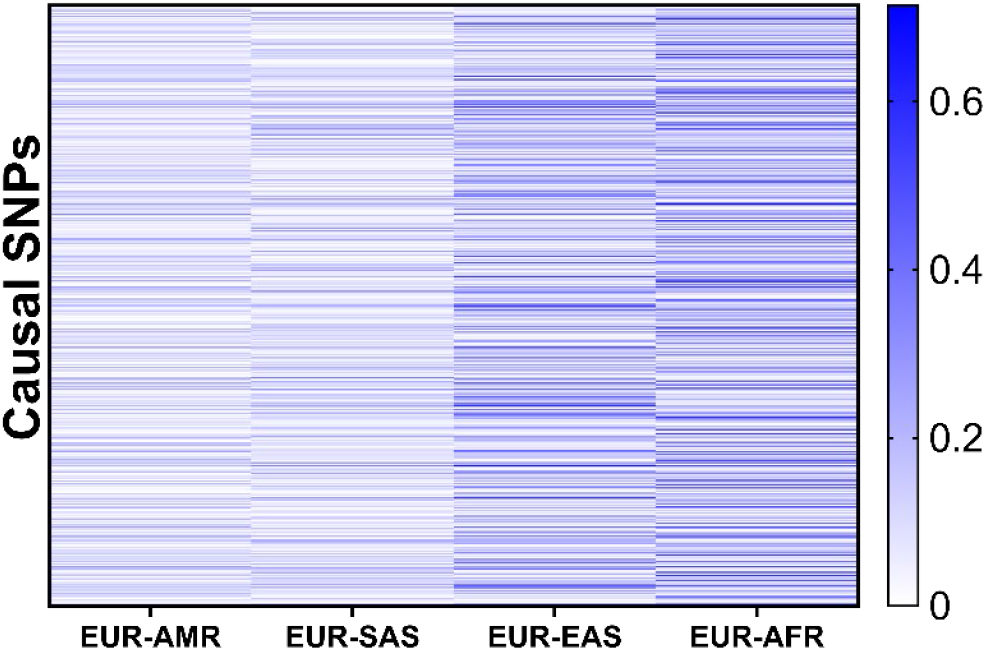
Allele frequency difference between EUR and four data-disadvantaged populations. The heatmap shows the absolute difference in minor allele frequencies between the EUR population and the data-disadvantaged populations (DDPs), AMR, SAS, EAS, and AFR. Each row represents an SNP. The value of each cell 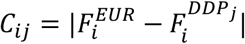, where 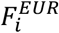 is the frequency of the minor allele of *i*^*th*^ SNP in the EUR population, and 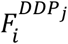 is the frequency of the same SNP allele in the *j*^*th*^ DDP.

From the multi-ancestral machine learning experiments on all the real and synthetic datasets (**Table 1, Fig. 2; Table 2, Fig. 4;** and **Table S2, Fig. S1**), we observe a consistent performance pattern that is characterized by four attributes: 1) Mixture learning generating model performance gap between the European population and the DDPs; 2) Independent learning generating model performance gap between the European population and the DDPs; 3) Low performance of naïve transfer learning, indicating poor generalizability of the model trained on EUR data to other ancestry groups; and 4) Transfer learning outperforming mixture learning, independent learning and naive transfer on the DDPs. The results on synthetic data show that the performance pattern is consistent for different DDPs and simulation parameters (**Table 2, Fig. 4;** and **Table S2, Fig. S1**). As expected, the heritability (*h*^2^) affects the prediction performance in all seven experiments. However, the performance pattern remains the same at different levels of heritability (*h*^2^ = 0.5 *and h*^2^ = 0.25). The parameter *ρ* is the correlation of the genetic effect sizes and is set as a function of genetic distance between ancestry groups (see the Methods section). Therefore, *ρ* represents the data distribution discrepancy between the EUR population and the DDPs. Our results show that the performance pattern is also robust against the variations of *ρ*. The overall performance comparison of the multi-ancestral machine learning schemes for the 14 synthetic datasets. For all the real and synthetic datasets, transfer learning provides the best performance on the DDPs.

**Fig. 4.**
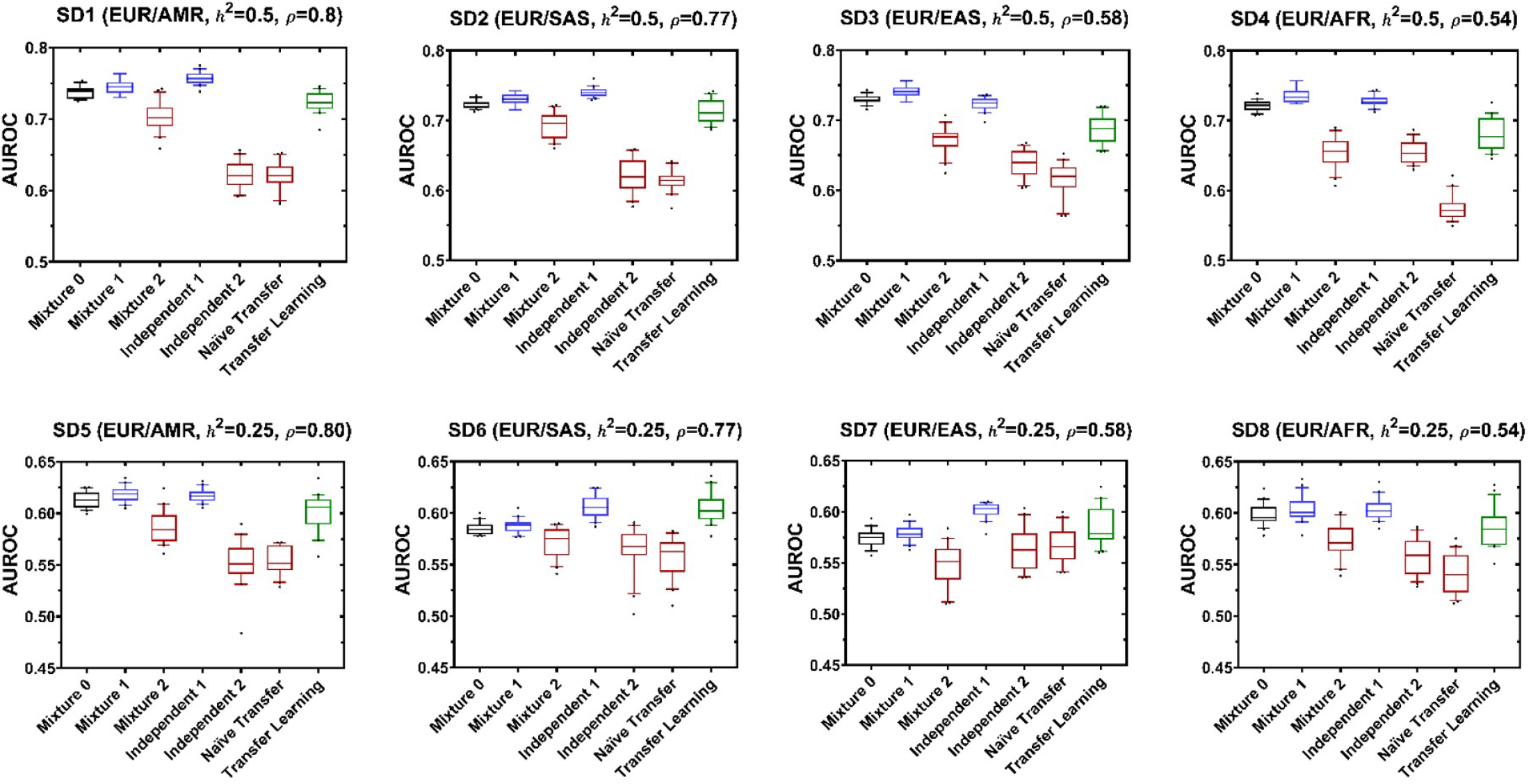
Comparison of multi-ancestral machine learning schemes on synthetic data. In each panel, the box plots show the AUROC values for the seven experiments (20 independent runs for each experiment). Box-plot elements are center line, median; box limits, 25 and 75 percentiles; whiskers, 10–90 percentiles; points, outliers.

## Discussion

Precision medicine is becoming increasingly dependent on high-performance machine learning models. Biomedical datasets are the essential foundation for the development of critical AI models for disease diagnostics, prognostics, and therapies. However, genomic data inequality severely impedes equitable precision medicine, posing a significant health risk to the data-disadvantaged populations. Currently, mixture learning is the most frequently used machine learning scheme for multi-ancestral data. The overall performance of the mixture learning model is mainly driven by its performance for the majority subpopulation in the dataset, which is usually the European ancestry group. The performance disparity gap between different subpopulations is often overlooked. We need a new multi-ancestral machine learning standard that requires machine learning models based on data from primarily European ancestry to be regularly tested for their validity, applicability, and accuracy in other ancestry groups.

Our study shows that transfer learning can improve disease prediction for data-disadvantaged populations, therefore, reduce the detrimental effects of genomic data inequality. The transfer learning scheme provides a *Pareto* improvement toward equitable machine learning for genomic medicine since it is not subject to the fairness-accuracy tradeoff, in contrast to the widely utilized constraint-based machine learning fairness approaches (**Fig.1**). A *Pareto* improvement is desired because it creates a new situation in which certain parties in a system are better off without negatively affecting other parties in the system^40^.

## Methods

### Data processing

The genotype, phenotype, and clinical data for lung cancer, prostate cancer, and Alzheimer’s disease were downloaded from the dbGaP database (phs001273.v3.p2, phs001391.v1.p1, phs000496.v1.p1, and phs000372.v1.p1)^41-43,46-50^. Genetic ancestries of the GWAS participants calculated using GRAF-pop^51^ were also downloaded from dbGaP.

We followed a standard quality-control process^52^ using plink^53^ (V1.9) to identify disease-associated single nucleotide polymorphism (SNPs) and remove outlier samples: 1) We filtered SNPs with a missing rate larger than 20% and then filtered samples with a missing SNP rate of higher than 20% and filtered samples with sex discrepancy; 2) We removed genetic marker SNPs with minor allele frequency (MAF) lower than 0.05 to reduce the size of feature set; 3) We removed the SNPs with a Hardy-Weinberg Equilibrium (HWE) p-value less than 10^−5^; 4) To eliminate data redundancy from Linkage Disequilibrium (LD), we used a sliding window of 50 SNPs and set the step length to 5 SNPs and set the LD cut-off coefficient to 0.2. The lung cancer feature set for our machine learning experiments consists of the top 500 SNPs with the smallest p-values from the association analysis, age, sex, and smoking status. The prostate cancer feature set consists of the top 1000 SNPs from the association analysis, age, and family history as the input variables.

We downloaded the lists of SNPs used in the construction of 16 polygenic scores for Alzheimer’s disease from the Polygenic Score (PGS) Catalog (https://www.pgscatalog.org/trait/MONDO_0004975/), including PGS000025, PGS000026, PGS000334, PGS000779, PGS000811, PGS000812, PGS000823, PGS000876, PGS000898, PGS000945, PGS001348, PGS001349, PGS001775, PGS001828, PGS002280, and PGS002731. The Polygenic Scores derived from the dbGaP datasets used in this study were excluded to avoid information leakage in feature selection^54,55^. We assembled a list of 22 SNPs that are used by both the PGS and the dbGaP datasets for Alzheimer’s disease. The Alzheimer’s disease feature set consists of the 22 SNPs, sex, and APOE (allele value for the Apolipoprotein E gene).

The processed datasets for our machine learning experiments are summarized in **Table S1**. We split each dataset into three parts: training set, validation set, and testing set, each comprised of 80%, 10%, and 10% of the participants, respectively, stratified by ancestry and class label (case/control). We randomly split the validation set and testing set with 20 different random seeds. The ancestry and case/control proportions were kept the same for all three sets in each partition. The association analysis for the training set was conducted using the additive logistic regression model from the PLINK software. We used a p-value threshold (p<10^−3^) to select the top SNPs. The ANOVA F-value for each SNP was calculated for the training samples to select the input features for machine learning. We selected the top 500 SNPs for lung cancer and 1000 SNPs for prostate cancer. The feature mask, ANOVA F-value, and p-values were calculated using the *SelectKBest* function in the Python-based scikit-learn (sklearn) machine learning software library.

### Data simulation

The simulated genotype data of the five ancestry groups^32^ were downloaded from the Harvard Dataverse^45^. We developed a statistical model to generate the disease case/control status. Each synthetic dataset *D* is a combination of two subsets, 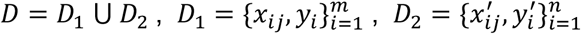, where *D*_1_ represents the European group, *D*_2_ represents the DDP group, *m* and *n* are the numbers of samples in the two groups, *x*_*ij*_ is the *j*^*th*^ feature of sample *i* in the European group, 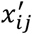 is the *j*^*th*^ feature of sample *i* in the DDP group, *y*_*i*_ is the case/control label of sample *i* in the European group, 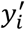 is the case/control label of sample *i* in the DDP group. The feature matrix *x*_*ij*_ and 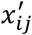 were sampled from the multi-ancestral genotype dataset, which contains simulated whole-genome genotype data for 120,000 individuals of each of the five ancestries, African (AFR), Admixed American (AMR), East Asian (EAS), European (EUR), and South Asian (SAS). We applied a quality-control process for each ancestry group to remove the SNPs with MAF<0.05 or HWE p-value < 10^−4^. We randomly selected 500 SNPs as the causal SNPs, 10,000 EUR individuals, and 2,000 individuals from each of the four non-EUR populations. We assembled four datasets, and each contains the genotype data of the selected causal SNPs for the 10,000 EUR individuals and 2,000 individuals of one of the non-EUR populations. The label of the *i*^*th*^ EUR individual is generated using the function 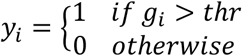, where 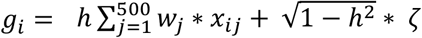 is the median value of {*g*_*i*_}, *w*_*j*_ is the effect size of the *j*^*th*^ SNP, *ζ* is sampled from a standard normal distribution with a truncated domain [-1,1], and *h*^2^ represents heritability. Similarly, The label of the *i*^*th*^ non-EUR individual is generated using the function 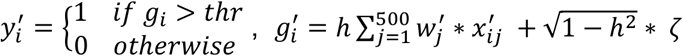, where *thr′* is the median value of 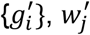 is the effect size of the *j*^*th*^ SNP. The SNP genetic effect vector for the European group *W* = [*w*_1_, *w*_2_, *w*_3_, … *w*_*n*_] was sampled from a random uniform distribution over [0,1]. 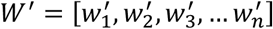 is generated using the following equation: 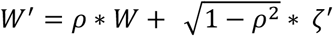, where *ρ* is the correlation of the genetic effect sizes between EUR and the DDP, *ζ*^′^ is sampled from a standard normal distribution with a truncated domain [-1,1]. We ensembled synthetic datasets that contain data from EUR and one of the four DDPs: {EUR, AFR}, {EUR, AMR}, {EUR, EAS}, {EUR, SAS}. We used 14 different combinations of the values of *h*^2^ and *ρ* to generate the synthetic datasets (**Table 2** and **Table S2**). We generated synthetic data at the two different levels of heritability (*h*^2^ = 0.5 *and h*^2^=0.25). The parameter *ρ* is a function of the genetic distance between the populations. The genetic distance between EUR and the *j*^*th*^ DDP is 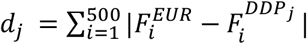, where 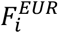 is the frequency of the minor allele of *i*^*th*^ causal SNP in the EUR population, and 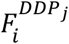 is the frequency of the same allele in the *j*^*th*^ DDP. The genetic distances are *d*_0_ = 42.7 (EUR - AMR), *d*_1_ = 46.1 (EUR - SAS), *d*_2_ = 80.6 (EUR - EAS), *d*_3_ = 94.1 (EUR - AFR). We used the function 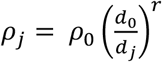 to generate values for parameter *ρ*, where *ρ*_0_ = 0.8. We set to *r* = 0.5 to generate the *ρ* values for synthetic datasets 1-8 (**Table 2**) and set *r* = 1.0 to generate the *ρ* values for synthetic datasets 9-16 (**Table S2**, SD9 and SD13 are the same as SD1 and SD5).

### Deep neural network and transfer learning

We used the Keras (https://keras.io/), and Tensorflow (https://www.tensorflow.org/) software packages to build the deep neural network (DNN) models. We used a pyramid architecture^56^ with four layers: an input layer with K nodes for the input features of genetic and other risk factors (if there is any), where the input feature number K=503 for the lung cancer dataset, including 500 SNPs, age, sex, and smoking status, K=1002 for the prostate cancer dataset, including 1,000 SNPs, age, and family history, K=24 for the Alzheimer’s disease datasets, including 22 SNPs, sex, and APOE (allele value for the Apolipoprotein E gene), and K=500 for the synthetic datasets, including 500 causal SNPs; two hidden layers including a fully connected layer with 100 nodes followed by a dropout layer^57^; and a logistic regression output layer. We used the stochastic gradient descent (SGD) algorithm with a learning rate of 0.25 to minimize a loss function consisting of a binary cross-entropy term and two regularization terms: 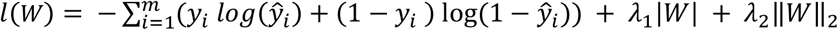, where *y*_*i*_ is the observed control/case label for individual *i*, ŷ_*i*_ is the predicted label for individual *i*, and *W* represents the weights in the DNN. We applied the ReLU activation function *f*(*x*) = *max*(0, *x*) to the output of the hidden layer to avoid the gradient vanish problem. For each drop out layer, we set the dropout probability *p* = 0.5 to randomly omit half of the weights during the training to reduce the collinearity between feature detectors. We split the data into multiple mini-batches (batch size = 32) for training to speed up the computation and improve the model prediction performance. We set the maximum number of iterations at 200 and applied the Nesterov momemtum^58^ method (with momentum=0.9 for each DNN model) to avoid premature stopping. We set the learning rate decay factor at 0.003 to avoid non-convergence during training. We also used early stopping with a patience value of 200 iterations to monitor the validation accuracy during model fitting. The two regularization terms λ_1_ and λ_2_ were set at 0.001.

In the transfer learning scheme, knowledge learned from the source domain, where training data is abundant, is transferred to assist the learning task for the target domain, where training data is inadequate^59-66^. The transferred knowledge is not the explicit knowledge expressed using mathematical or natural languages but is encoded in the machine learning model, for example, the structure and parameters of a neural network. We used the European ancestry population as the source domain and a DDP as the target domain. We adopted a supervised fine-tuning algorithm for transfer learning. We first pretrained a DNN model using the source domain data: *M* ∼ *f*(*Y*_*source*_|*X*_*source*_), wehre *Y*_*source*_ is the class label of each individual in the European group, *X*_*source*_ is the input feature data of the European group. We trained the model using the parameters described above. After the pretraining, we fine-tuned the DNN model with the backpropagation method using the target domain data: *M*^′^ = *fine*_*tuning*(*M* | *Y*_*Target*_, *X*_*Target*_), where *M*^′^ is the final model. In the fine-tuning, the learning rate was set at 0.25, and the batch size was set at 32, as the model has been partially fitted and the target domain dataset is smaller.

We performed 20 independent runs for each experiment. Each run uses the same training set but a different random splitting of validation and testing sets. We used the Area Under ROC Curve (AUROC)^44^ to evaluate machine learning model performance. We used the one-sided paired t-test to calculate the p-values for the assessment of statistical significance in the machine learning model performance comparisons.

## Supplementary information

**Table S1:** The datasets used for machine learning experiments.

| **Lung Cancer** |  |  |
| --- | --- | --- |
| European | Case | 9675 |
|  | Control | 8107 |
| East Asian | Case | 1363 |
|  | Control | 801 |
| **Prostate Cancer** |  |  |
| European | Case | 28385 |
|  | Control | 19829 |
| African American | Case | 1418 |
|  | Control | 2021 |
| **Alzheimer's Disease** |  |  |
| European | Case | 3632 |
|  | Control | 1588 |
| Latin American | Case | 951 |
|  | Control | 1078 |
| African American | Case | 682 |
|  | Control | 821 |

**Table S2.** Multi-ancestral machine learning experiments on synthetic data.

| Synthetic Datasets |  |  |  | AUROC |  |  |  |  |  |  |
| --- | --- | --- | --- | --- | --- | --- | --- | --- | --- | --- |
| ID | DDP | $h^2$ | $\rho$ | Mixture 0 | Mixture 1 | Mixture 2 | Independent 1 | Independent 2 | Naive Transfer | Transfer Learning |
| SD9* | AMR | 0.50 | 0.80 | 0.74 | 0.75 | 0.70 | 0.76 | 0.62 | 0.62 | 0.72 |
| SD10 | SAS | 0.50 | 0.74 | 0.72 | 0.72 | 0.70 | 0.73 | 0.64 | 0.61 | 0.72 |
| SD11 | EAS | 0.50 | 0.42 | 0.66 | 0.67 | 0.59 | 0.72 | 0.63 | 0.59 | 0.66 |
| SD12 | AFR | 0.50 | 0.36 | 0.65 | 0.68 | 0.53 | 0.73 | 0.63 | 0.55 | 0.66 |
| SD13* | AMR | 0.25 | 0.80 | 0.61 | 0.62 | 0.59 | 0.62 | 0.55 | 0.55 | 0.60 |
| SD14 | SAS | 0.25 | 0.74 | 0.58 | 0.59 | 0.57 | 0.60 | 0.57 | 0.55 | 0.62 |
| SD15 | EAS | 0.25 | 0.42 | 0.58 | 0.59 | 0.54 | 0.61 | 0.55 | 0.55 | 0.57 |
| SD16 | AFR | 0.25 | 0.36 | 0.60 | 0.60 | 0.57 | 0.61 | 0.57 | 0.53 | 0.58 |

**AUROC:** The mean Area under ROC Curve values from 20 independent runs for each experiment; **SD:** Synthetic data; **EUR:** European population; **DDP:** Data-disadvantaged population; **AFR:** African; **AMR:** Admixed American; **EAS:** East Asian; **SAS:** South Asian. \*SD9 and SD13 are the same as SD1 and SD5 and are included here for comparison.

**Fig. S1.**
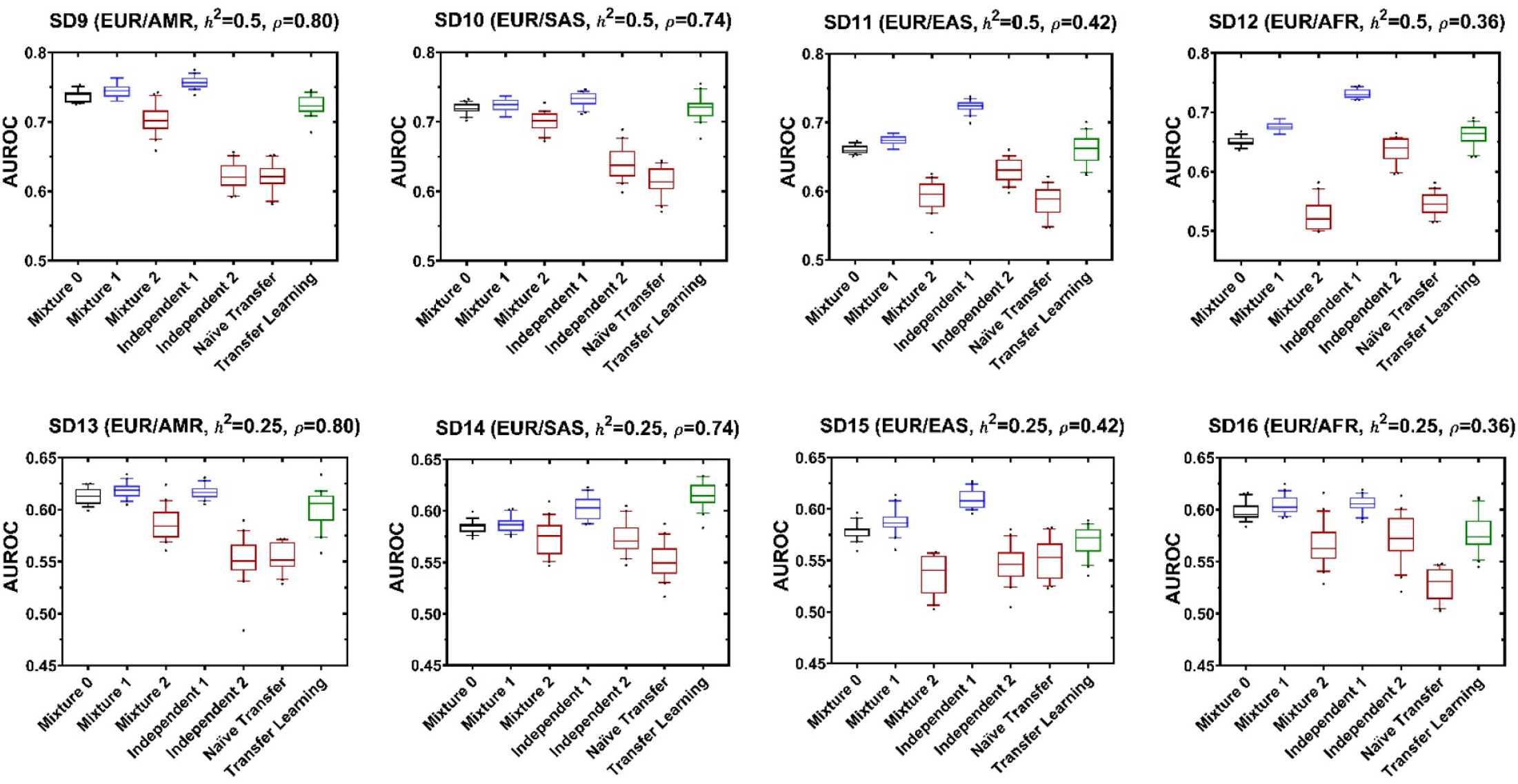
Comparison of multi-ancestral machine learning schemes using synthetic data. The box plots in each panel show the AUROC values for the seven experiments (20 independent runs for each experiment). Box-plot elements are center line, median; box limits, 25 and 75 percentiles; whiskers, 10–90 percentiles; points, outliers. SD9 and SD13 are the same as SD1 and SD5 and are included here for comparison.

## References

1. Mills, M.C. & Rahal, C. The GWAS Diversity Monitor tracks diversity by disease in real time. Nature Genetics 52, 242–243 (2020).

2. GWAS Diversity Monitor. (https://gwasdiversitymonitor.com).

3. Sirugo, G., Williams, S.M. & Tishkoff, S.A. The Missing Diversity in Human Genetic Studies. Cell 177, 26–31 (2019).

4. Gurdasani, D., Barroso, I., Zeggini, E. & Sandhu, M.S. Genomics of disease risk in globally diverse populations. Nature Reviews Genetics 20, 520–535 (2019).

5. Guerrero, S. et al. Analysis of Racial/Ethnic Representation in Select Basic and Applied Cancer Research Studies. Scientific Reports 8, 13978 (2018).

6. Martin, A.R. et al. Clinical use of current polygenic risk scores may exacerbate health disparities. Nature genetics 51, 584–591 (2019).

7. Bien, S.A. et al. The Future of Genomic Studies Must Be Globally Representative: Perspectives from PAGE. Annual review of genomics and human genetics (2019).

8. Gao, Y. & Cui, Y. Deep transfer learning for reducing health care disparities arising from biomedical data inequality. Nature Communications 11, 5131 (2020).

9. Torkamani, A., Wineinger, N.E. & Topol, E.J. The personal and clinical utility of polygenic risk scores. Nature Reviews Genetics 19, 581–590 (2018).

10. Lambert, S.A., Abraham, G. & Inouye, M. Towards clinical utility of polygenic risk scores. Human Molecular Genetics 28, R133–R142 (2019).

11. Lewis, C.M. & Vassos, E. Polygenic risk scores: from research tools to clinical instruments. Genome Medicine 12, 44 (2020).

12. Choi, S.W., Mak, T.S.-H. & O’Reilly, P.F. Tutorial: a guide to performing polygenic risk score analyses. Nature Protocols 15, 2759–2772 (2020).

13. Wray, N.R. et al. From Basic Science to Clinical Application of Polygenic Risk Scores: A Primer. JAMA Psychiatry 78, 101–109 (2021).

14. Polygenic Risk Score Task Force of the International Common Disease, A. Responsible use of polygenic risk scores in the clinic: potential benefits, risks and gaps. Nat Med 27, 1876–1884 (2021).

15. Kullo, I.J. et al. Polygenic scores in biomedical research. Nat Rev Genet 23, 524–532 (2022).

16. Ma, Y. & Zhou, X. Genetic prediction of complex traits with polygenic scores: a statistical review. Trends in Genetics 37, 995–1011 (2021).

17. Dai, Z., Long, N. & Huang, W. Influence of Genetic Interactions on Polygenic Prediction. G3: Genes|Genomes|Genetics 10, 109–115 (2020).

18. Uddin, S., Khan, A., Hossain, M.E. & Moni, M.A. Comparing different supervised machine learning algorithms for disease prediction. BMC medical informatics and decision making 19, 1–16 (2019).

19. Ho, D.S.W., Schierding, W., Wake, M., Saffery, R. & O’Sullivan, J. Machine Learning SNP Based Prediction for Precision Medicine. Frontiers in Genetics 10(2019).

20. Gao, Y. & Cui, Y. Clinical time-to-event prediction enhanced by incorporating compatible related outcomes. PLOS Digital Health 1(2022).

21. Badré, A., Zhang, L., Muchero, W., Reynolds, J.C. & Pan, C. Deep neural network improves the estimation of polygenic risk scores for breast cancer. Journal of Human Genetics (2020).

22. Elgart, M. et al. Non-linear machine learning models incorporating SNPs and PRS improve polygenic prediction in diverse human populations. Communications Biology 5, 856 (2022).

23. Leist, A.K. et al. Mapping of machine learning approaches for description, prediction, and causal inference in the social and health sciences. Science Advances 8, eabk1942 (2022).

24. Martin, A.R. et al. Human Demographic History Impacts Genetic Risk Prediction across Diverse Populations. The American Journal of Human Genetics 100, 635–649 (2017).

25. Duncan, L. et al. Analysis of polygenic risk score usage and performance in diverse human populations. Nature Communications 10, 3328 (2019).

26. Chen, M.-H. et al. Trans-ethnic and Ancestry-Specific Blood-Cell Genetics in 746,667 Individuals from 5 Global Populations. Cell 182, 1198-1213.e14 (2020).

27. Zhou, W. et al. Global Biobank Meta-analysis Initiative: powering genetic discovery across human diseases. medRxiv, 2021.11.19.21266436 (2021).

28. Wang, Y. et al. Theoretical and empirical quantification of the accuracy of polygenic scores in ancestry divergent populations. Nat Commun 11, 3865 (2020).

29. Prive, F. et al. Portability of 245 polygenic scores when derived from the UK Biobank and applied to 9 ancestry groups from the same cohort. Am J Hum Genet 109, 12–23 (2022).

30. Ruan, Y. et al. Improving polygenic prediction in ancestrally diverse populations. Nat Genet 54, 573–580 (2022).

31. Cai, M. et al. A unified framework for cross-population trait prediction by leveraging the genetic correlation of polygenic traits. Am J Hum Genet 108, 632–655 (2021).

32. Zhang, H. et al. Novel Methods for Multi-ancestry Polygenic Prediction and their Evaluations in 3.7 Million Individuals of Diverse Ancestry. bioRxiv, 2022.03.24.485519 (2022).

33. Coram, M.A., Fang, H., Candille, S.I., Assimes, T.L. & Tang, H. Leveraging Multi-ethnic Evidence for Risk Assessment of Quantitative Traits in Minority Populations. Am J Hum Genet 101, 218–226 (2017).

34. Xiao, J. et al. XPXP: improving polygenic prediction by cross-population and cross-phenotype analysis. Bioinformatics 38, 1947–1955 (2022).

35. Weissbrod, O. et al. Leveraging fine-mapping and multipopulation training data to improve cross-population polygenic risk scores. Nat Genet 54, 450–458 (2022).

36. Gao, Y. & Cui, Y. Multi-ethnic Survival Analysis: Transfer Learning with Cox Neural Networks. in Proceedings of AAAI Spring Symposium on Survival Prediction - Algorithms, Challenges, and Applications 2021 Vol. 146 (eds Russ, G., Neeraj, K., Thomas Alexander, G. & Mihaela van der, S.) 252--257 (PMLR, Proceedings of Machine Learning Research, 2021).

37. Mehrabi, N., Morstatter, F., Saxena, N., Lerman, K. & Galstyan, A. A Survey on Bias and Fairness in Machine Learning. ACM Computing Surveys 54, 1–35 (2021).

38. Zhao, H. & Gordon, G. Inherent tradeoffs in learning fair representations. Advances in neural information processing systems 32(2019).

39. Menon, A.K. & Williamson, R.C. The cost of fairness in binary classification. in Conference on Fairness, Accountability and Transparency 107-118 (PMLR, 2018).

40. Chatterjee, D.K. Encyclopedia of Global Justice, (Springer Science & Business Media, 2011).

41. McKay, J.D. et al. Large-scale association analysis identifies new lung cancer susceptibility loci and heterogeneity in genetic susceptibility across histological subtypes. Nat Genet 49, 1126–1132 (2017).

42. Amos, C.I. et al. The OncoArray Consortium: A Network for Understanding the Genetic Architecture of Common Cancers. Cancer Epidemiology Biomarkers & Prevention 26, 126–135 (2017).

43. Timofeeva, M.N. et al. Influence of common genetic variation on lung cancer risk: meta-analysis of 14 900 cases and 29 485 controls. Human molecular genetics 21, 4980–4995 (2012).

44. Fawcett, T. An introduction to ROC analysis. Pattern Recognition Letters 27, 861–874 (2006).

45. Harvard Dataverse. (https://doi.org/10.7910/DVN/COXHAP).

46. Reitz, C. et al. A Summary Risk Score for the Prediction of Alzheimer Disease in Elderly Persons. Archives of Neurology 67, 835–841 (2010).

47. Lee, J.H. et al. Identification of Novel Loci for Alzheimer Disease and Replication of CLU, PICALM, and BIN1 in Caribbean Hispanic Individuals. Archives of Neurology 68, 320–328 (2011).

48. Naj, A.C. et al. Common variants at MS4A4/MS4A6E, CD2AP, CD33 and EPHA1 are associated with late-onset Alzheimer’s disease. Nature Genetics 43, 436–441 (2011).

49. Jun, G. et al. Meta-analysis Confirms CR1, CLU, and PICALM as Alzheimer Disease Risk Loci and Reveals Interactions With APOE Genotypes. Archives of Neurology 67, 1473–1484 (2010).

50. The database of Genotypes and Phenotypes. (https://www.ncbi.nlm.nih.gov/gap/).

51. Jin, Y., Schaffer, A.A., Feolo, M., Holmes, J.B. & Kattman, B.L. GRAF-pop: a fast distance-based method to infer subject ancestry from multiple genotype datasets without principal components analysis. G3: Genes, Genomes, Genetics 9, 2447–2461 (2019).

52. Marees, A.T. et al. A tutorial on conducting genome-wide association studies: Quality control and statistical analysis. International journal of methods in psychiatric research 27, e1608 (2018).

53. Purcell, S. et al. PLINK: a tool set for whole-genome association and population-based linkage analyses. The American journal of human genetics 81, 559–575 (2007).

54. Refaeilzadeh, P., Tang, L. & Liu, H. On comparison of feature selection algorithms. in Proceedings of AAAI workshop on evaluation methods for machine learning II Vol. 3 5 (AAAI Press Vancouver, 2007).

55. Molla, M., Waddell, M., Page, D. & Shavlik, J. Using Machine Learning to Design and Interpret Gene-Expression Microarrays. AI Magazine 25, 23 (2004).

56. Phung, S.L. & Bouzerdoum, A. A pyramidal neural network for visual pattern recognition. IEEE transactions on neural networks 18, 329–343 (2007).

57. Srivastava, N., Hinton, G., Krizhevsky, A., Sutskever, I. & Salakhutdinov, R. Dropout: a simple way to prevent neural networks from overfitting. The journal of machine learning research 15, 1929–1958 (2014).

58. Sutskever, I., Martens, J., Dahl, G. & Hinton, G. On the importance of initialization and momentum in deep learning. in International conference on machine learning 1139-1147 (2013).

59. Yang, Q., Zhang, Y., Dai, W. & Pan, S.J. Transfer learning, (Cambridge University Press, 2020).

60. Pan, S.J. & Yang, Q. A survey on transfer learning. IEEE Transactions on knowledge and data engineering 22, 1345–1359 (2010).

61. Tan, C. et al. A survey on deep transfer learning. in International Conference on Artificial Neural Networks 270–279 (Springer, 2018).

62. Weiss, K., Khoshgoftaar, T.M. & Wang, D. A survey of transfer learning. Journal of Big data 3, 9 (2016).

63. Taroni, J.N. et al. MultiPLIER: a transfer learning framework for transcriptomics reveals systemic features of rare disease. Cell systems 8, 380–394 (2019).

64. Wang, J. et al. Data denoising with transfer learning in single-cell transcriptomics. Nature Methods 16, 875–878 (2019).

65. Sevakula, R.K., Singh, V., Verma, N.K., Kumar, C. & Cui, Y. Transfer Learning for Molecular Cancer Classification Using Deep Neural Networks. IEEE/ACM Transactions on Computational Biology and Bioinformatics 16, 2089–2100 (2019).

66. Ebbehoj, A., Thunbo, M.Ø., Andersen, O.E., Glindtvad, M.V. & Hulman, A. Transfer learning for non-image data in clinical research: A scoping review. PLOS Digital Health 1, e0000014 (2022).

